# Flex-It: A global standardised genotyping framework for *Shigella flexneri*

**DOI:** 10.64898/2026.04.17.719127

**Authors:** Jane Hawkey, Carolina S Nodari, Zamin Iqbal, Martin Hunt, Ryan R Wick, Charlotte E Chong, Claire Jenkins, Benjamin P Howden, Kathryn E Holt, François-Xavier Weill, Kate S Baker, Danielle J Ingle

**Author notes:** **Corresponding authors** Jane Hawkey **E:** Danielle J Ingle **E:**.

## Abstract

*Shigella flexneri* is the leading causative agent of shigellosis globally. The public health threat posed by *S. flexneri* is compounded by its emergence as a sexually transmissible infection, importance of international travel in driving dissemination, and the increasing prevalence of antimicrobial resistance (AMR). A rapid and robust computational method is needed to enhance genomic surveillance and systematically explore features of the population structure of this WHO priority pathogen, which is scalable and readily implementable across jurisdictions, particularly as vaccine development efforts are underway. Here, we present Flex-It, a genomic framework and genotyping scheme implemented in Mykrobe for *S. flexneri* serotypes 1-5, X & Y, compatible with previous approaches used to describe *S. flexneri’s* population structure. To develop Flex-It, we curated a retrospective dataset of 5,819 publicly available *S. flexneri* genomes. We characterised the global population structure for *S. flexneri*, exploring geographical and temporal traits, and showed the granular diversity of AMR and serotype profiles. We applied Flex-It to >13,000 genomes routinely generated by public health laboratories from Australia, the UK and the USA across a ten-year period. We found significant genotype diversity in all three locations, with the emergence of genotypes with converged resistance to all major drugs currently used for treatment. Flex-It provides an open-source, novel genotyping method that rapidly characterises *S. flexneri* and its ciprofloxacin resistance determinants in <1 minute from both short and long whole-genome sequencing reads. Flex-It provides the community with a standardised nomenclature to monitor the emergence and spread of *S. flexneri* lineages.

## Introduction

*Shigella* is the causative agent of shigellosis, which spreads via the faecal-oral route and is the leading bacterial cause of diarrhoeal disease in children under five years of age in low-middle income countries (LMICs)^1,2^. Clinical manifestations of shigellosis range from watery diarrhoea to dysentery, and recurrent infections can result in long-term poor health outcomes such as stunting and impaired cognitive function^1,3,4^. *Shigella flexneri* is one of four serogroups (formerly species) within the ‘*Shigella’* genus (the other three serogroups are *S. sonnei, S. boydii* and *S. dysenteriae*)^5^ and causes approximately two thirds of the paediatric disease burden in LMICs^1,6,7^. Phylogenetically, these serogroups have emerged from within the *Escherichia coli* species, driven by acquisition of the virulence plasmid and gene loss events^5,8^. For historical and clinical reasons, they are still often classified separately from *E. coli,* although not all *Shigella* serogroups are monophyletic (**Supplementary Figure 1)**. Notably, *Shigella* are considered priority pathogens by the WHO due to both pandemic potential and increasing antimicrobial resistance (AMR), in particular resistance to recommended empiric treatments including fluoroquinolones, third-generation cephalosporins (3GCs), and macrolides^9–11^.

Although infections caused by *S. flexneri* are disproportionately experienced LMICs, particularly in children^1^, *S. flexneri* is second only to *S. sonnei* as a cause of shigellosis^12,13^. Historically considered a contaminated food or water borne pathogen in LMICs and associated with returning travellers in high-income countries (HICs), the epidemiology of *S. flexneri* has shifted in the past decade^13–16^. Transmission has increased among at-risk groups in HICs, where *S. flexneri* is now an endemic sexually transmissible infection (STI) among men who have sex with men (MSM)^15,17–19^. This change in epidemiology is thought to be associated with changing behaviours and the rise of AMR^17,20–24^. Multi-drug resistance (MDR, defined as resistance to three or more drug classes) is increasing in *S. flexneri*, mediated by chromosomally integrating elements and large plasmids that carry AMR mechanisms to key therapeutics including azithromycin and trimethoprim-sulfamethoxazole^14,24,25^. More recently, resistance to ciprofloxacin and 3GCs including ceftriaxone have also been detected, giving rise to extensively drug-resistant (XDR) strains (defined as resistance to all but two classes of antimicrobials^17,26^, and usually refers to triple-resistant due to resistance to fluoroquinolones, third-generation cephalosporins, and azithromycin in *Shigella*^16,17,24,27–30^). Further, previous studies have found that population expansions of XDR *S. flexneri* strains are emerging in both HICs and LMICs^20,21,23,28,31^.

Discriminative identification among *S. flexneri* subtypes has traditionally been achieved with serotyping. Unlike the monophyletic and monoserotypic *S. sonnei, S. flexneri* has significantly greater genetic diversity. It has arisen from multiple lineages within *E. coli* and has several distinct serotypes^19^. The majority of *S. flexneri* belong to clonal complex (CC) 245, and consist of serotypes 1–5, X and Y. This group accounts for ∼80% of *S. flexneri* recovered in epidemiological surveillance^1,32,33^. In contrast, serotype 6 belongs to a distinct *S. flexneri* clade that represents an independently evolved genomic lineage (CC145) of *E. coli* (see **Supplementary Figure 1**)^19,32^. In the genomic era, *S. flexneri* serotypes 1–5, X and Y have been further defined by Connor *et al*^19^ into seven phylogroups (PGs) that capture its high-level population diversity^19^. Each PG has been shown to comprise multiple serotypes, some of which are multiphyletic, which complicates surveillance efforts relying on serotype^19,34^. Further, *S. flexneri* serotype switching events have been shown to occur in different PGs and over short timespans, which has implications for current efforts to develop and implement a *Shigella* vaccine^34^. Hence, robust genomic subtyping methods would offer a highly discriminative, complementary surveillance mechanism for this important pathogen.

Tracking the emergence and spread of AMR using genomics in *S. flexneri* remains a significant challenge. The analytical frameworks used for genomic epidemiology and public health surveillance of *S. flexneri* vary widely between settings^35^. Serotypes of *S. flexneri* often form the initial basis for public health investigations as this has been the traditional lens for surveillance. For example, emerging drug-resistant lineages have often been associated with subserotypes 2a, 3a and more recently 1b^17,24,36^. The seven PGs defined in 2015^19^ remain in use, although PG identification requires reference isolates and high-resolution phylogenetic analyses. More recently, core-genome MLST (cgMLST) approaches have been used, largely using the *E. coli* scheme developed and implemented in EnteroBase with the HierCC classifications^37^. As this scheme was developed for the broader *E. coli* species, it was not specifically designed to capture the unique population structure within *Shigella* and is not widely reported as part of *Shigella* investigations. Some public health agencies have developed their own in-house methods for genomic investigation of *S. flexneri*, such as ‘SNP-address’ or using manual curation of high-resolution phylogenetic analyses, but these rely on in-house databases, precluding translation across jurisdictions^13,36^. For *S. sonnei*, this has been addressed through a single nucleotide polymorphism (SNP)-based genotyping scheme that was introduced in 2021 and has been widely adopted; the same approach has also been successfully implemented for *Salmonella enterica* serovars Typhi and Paratyphi B^38–40^.

To address this remaining surveillance need for a high priority pathogen we present here Flex-It, a genomic framework and genotyping scheme. Flex-It is implemented in Mykrobe, an open-source *k*-mer-based software tool, that identifies genotype and ciprofloxacin resistance determinants directly from either short (Illumina) or long (Oxford Nanopore) whole-genome sequencing reads in <1 minute for *S. flexneri* serotypes 1-5, X & Y. The scheme is compatible with previous approaches used to describe *S. flexneri’s* population structure, including PGs and serotyping. To develop Flex-It, we curated a retrospective dataset of 5,819 publicly available *S. flexneri* genomes, characterising their global population structure and exploring geographical, temporal, and accessory genome traits. We then applied Flex-It to >13,000 routinely generated genomes from public health laboratories in three HICs across a ten-year period (2015-2024).

We demonstrate genotype diversity in all three locations, with emergence of genotypes converging on detection of AMR mechanism to all major drugs currently recommended for treatment. Flex-It provides an open-source, novel genotyping method to rapidly and robustly identify *S. flexneri* directly from reads with a standardised nomenclature that demonstrably enhances genomic surveillance across borders.

## Results

### *S. flexneri* diversity and distribution

The retrospective dataset consisted of 5,819 genomes from 23 previously published studies (see **Methods, Table 1**), spanning >100 years, with the earliest sample collected in 1913 to 2022, representing 97 countries and all inhabited continents (**Figure 1**). Serotype 6 genomes were excluded from the dataset as they represent a unique monophyletic lineage elsewhere in the phylogeny (**Supplementary Figure 1**). We defined nine PGs in the *S. flexneri* population structure (**Figure 1a**), adding two additional PGs to the original seven defined by Connor *et al*^19^. As per Yassine *et al*^41^, we used the cgMLST-based Enterobase HeirCC HC1100 groupings to define PGs, except for PGs 1 and 3 (see **Methods** for details). As all PGs were required to be monophyletic for developing the genotyping scheme; the paraphyletic PG4 was split into two separate PGs, PG4 and PG8. We additionally defined a new PG, PG9, which was not found in the original Connor *et al*^19^ study. PG9 represents a high-level lineage previously identified at the cgMLST HC1100 level by Yassine *et al*^41^, and is made up exclusively of *S. boydii* serotype 22 genomes.

**Figure 1:**
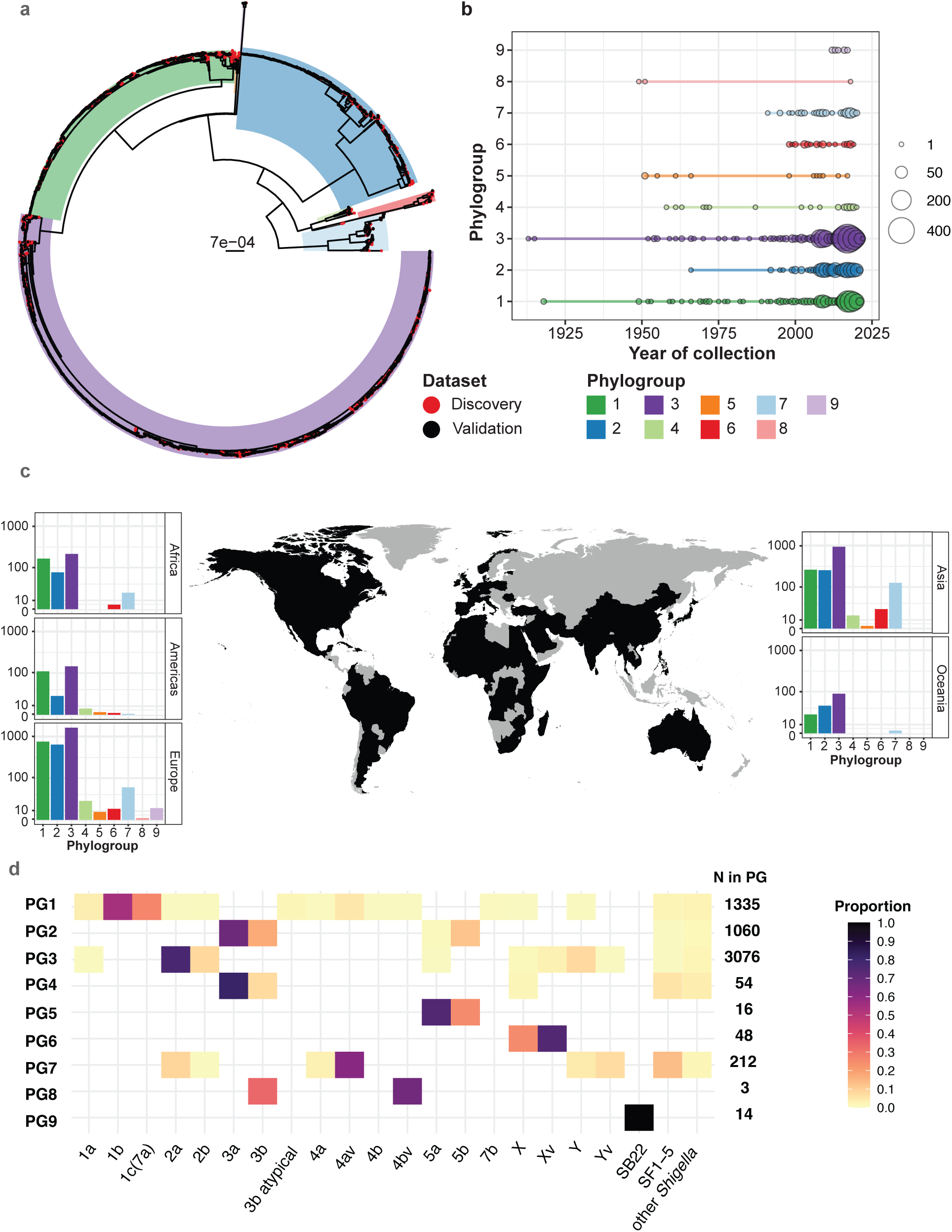
Overview o*f S. flexneri* dataset. **a**) Maximum-likelihood phylogeny of all 5,819 *S. flexneri* genomes. Phylogroups (PGs) are highlighted as per legend. Tips are coloured red if the genome belongs to the ‘discovery dataset’, black if it belongs to ‘validation dataset’. **b)** Distribution of year of collection by PG. Size of circle indicates the number of genomes present in that year. PGs are coloured as per legend and panel **a**. **c**) Countries where genomes were collected are indicated in black. Inset bar plots show the number of genomes per PG by world region. **d**) Proportion of genomes carrying a serotype, by PG. Coloured boxes show the proportion of genomes (as per legend) within PG carrying a particular serotype, predicted with ShigaPass. Serotype variants within serotypes 1-5, X & Y are given. SB22 indicates *S. boydii* serotype 22; SF1-5 indicates that ShigaPass determined this is an *S. flexneri* serotype but could be more specific; ‘other *Shigella’* indicates ShigaPass was unable to serotype more specifically than simply ‘*Shigella* spp.’. Total number of genomes per PG are listed in bold to the right of the heatmap.

**Table 1:** Summary of published studies included in historical dataset.

| Study | PMID | n | Year range |
| --- | --- | --- | --- |
| Albert 2021 *Gut Pathog* | PMID: 33743818 <sup>70</sup> | 1 | 2019 |
| Allen 2021 *Clin Microbiol Infect* | PMID: 32247893 <sup>71</sup> | 29 | 2010-2017 |
| Baker 2015 *Lancet Infect Dis* | PMID: 25936611 <sup>14</sup> | 258 | 1995-2014 |
| Baker 2018 *Nat Commun* | PMID: 29654279 <sup>15</sup> | 101 | 2008-2014 |
| Bardsley 2020 *J Clin Microbiol* | PMID: 31969425 <sup>57</sup> | 546 | 2015-2019 |
| Bengtsson 2021 *Nat Microbiol* | PMID: 35102306 <sup>34</sup> | 630 | 2007-2011 |
| Bernaquez 2021 *Microb Genom* | PMID: 34730485 <sup>72</sup> | 89 | 2017-2019 |
| Bogaerts 2021 *Microorganisms* | PMID: 33917583 <sup>73</sup> | 59 | 2013-2018 |
| Chattaway 2017 *J Clin Microbiol* | PMID: 27974538 <sup>74</sup> | 87 | 2005-2016 |
| Chattaway 2017 *Front Microbiol* | PMID: 28974944 <sup>75</sup> | 29 | 2015 |
| Connor 2015 *eLife* | PMID: 26238191 <sup>19</sup> | 333 | 1913-2011 |
| Ezernitchi 2019 *PLoS One* | PMID: 31626667 <sup>76</sup> | 15 | 2014-2016 |
| Ingle 2019 *Clin Infect Dis* | PMID: 30615105 <sup>13</sup> | 170 | 2016-2018 |
| Mitchell 2019 *Microb Genom* | PMID: 31682221 <sup>77</sup> | 913 | 2015-2017 |
| Moreno-Mingorance 2021<br>*Int J Antimicrob Agents* | PMID: 34157402 <sup>78</sup> | 44 | 2015-2019 |
| Parajuli 2020 *Genes* | PMID: 32899396 <sup>27</sup> | 69 | unknown |
| Rossi 2015 *Gut Pathog* | PMID: 26042182 <sup>79</sup> | 8 | 1954-2014 |
| Sethuvel 2020 *mSphere* | PMID: 33028681 <sup>80</sup> | 79 | 1990-2011 |
| Tansarli 2023 *Lancet Infect Dis* | PMID: 36731480 <sup>29</sup> | 103 | 2017-2022 |
| The 2021 *Commun Biol* | PMID: 33742111 <sup>16</sup> | 753 | unknown |
| Yassine 2022 *Nat Commun* | PMID: 35087053 <sup>41</sup> | 1425 | 2007-2021 |
| van den Beld 2020 *Front Microbiol* | PMID: 33193150 <sup>81</sup> | 75 | 2016-2017 |
| van den Beld 2018 *J Clin Microbiol* | PMID: 30021824 <sup>82</sup> | 3 | unknown |

Most genomes belonged to PG1, PG2 and PG3 (23%, 18% and 53%, respectively, **Figure 1a**), with PG1 and PG3 containing the oldest representatives (**Figure 1b**). As expected, 59.5% (n=3464/5819) of genomes have been collected since the advent and implementation of high-throughput WGS (defined here as 2015, **Figure 1b**). Most genomes originated from Europe (35%) or Asia (28%), likely due to large surveillance studies conducted in these regions, or integration of reported travel history to endemic regions for isolates collected from HICs (**Figure 1c, Supplementary Table 1**). PG1, PG2, PG3 and PG7 genomes originated in all geographical regions, in contrast to PG8 and PG9 which were only detected in Europe (**Figure 1c**, **Methods**). We performed *in silico* serotyping of all 5,819 genomes using ShigaPass^42^, which confirmed a diversity of serotypes within the nine PGs, with all PGs except PG9 harbouring ≥2 serotypes (**Figure 1d**). Our finding that individual serotypes were present across multiple PGs in the *S. flexneri* serogroup reinforces that the use of serotyping for tracking outbreaks of *S. flexneri*, and other surveillance aims, may lead to misleading results.

### Defining and validating the genotyping framework

Having curated and established a retrospective dataset that encapsulated the genetic diversity of *S. flexneri,* we split the 5,819 genomes into two groups, “discovery” and “validation”. The discovery dataset consisted of a phylogenetically diverse sampling of 1,999 genomes and the reference genome *S. flexneri* 2a str. 301 (total n=2,000; see **Methods**). This discovery dataset was used to define the genotyping scheme, which is a hierarchical scheme in the format of [PG].[lineage].[clade].[subclade], with additional groups below subclade defined for some genotypes (see **Methods**), similar to previous schemes for *S. sonnei, S.* Typhi, and *S.* Paratyphi B^38–40^. We retained the use of PG as the first index in the scheme to ensure backwards compatibility with the previously defined PGs^19^. For PGs 4-9, lineages were defined based on cgMLST HC400 thresholds, with clades and subclades defined by expert manual curation, incorporating tree structure, trends in epidemiological metadata, and previously identified epidemiological clusters (see **Methods**). For the larger phylogroups, PGs 1, 2 and 3, we defined lineages, clades and subclades using pairwise SNP distances (see **Methods, Supplementary Figure 2**), as per *S. sonnei*. However, unlike for *S. sonnei*, where the same cutoffs were applied across all lineages, separate cutoffs were defined for each PG, as a single *S. flexneri* PG contains a similar level of nucleotide diversity to the entirety of the *S. sonnei* species^34^. SNP cutoffs ranged between 800-850 for lineage, 400-450 for clade, and 140-210 for subclade, depending on PG (**Supplementary Figure 2**).

The final Flex-It genotyping scheme comprised of 119 marker SNPs, defining a total of 119 genotypes (**Figure 2**). **Supplementary Table 2** details the marker SNP for each genotype. Flex-It was implemented in the genotyping tool Mykrobe^43,44^, along with probes for detecting ciprofloxacin-resistance causing mutations in the quinolone resistance determining region (QRDR) genes *gyrA* and *parC*, as done previously for *S. sonnei*^40^. The scheme also includes a parser script for interpreting the Mykrobe output and scoring genotype calls as ‘strong’, ‘moderate’, or ‘weak’ according to the confidence Mykrobe places on each level of the genotype call (see **Methods**), as we have done previously for *S. sonnei* and *S.* Typhi^38,40^. While this provides information as to the confidence of the call, it does not replace the need for quality control to be undertaken by end-users. The Mykrobe implementation of Flex-It can be found at https://github.com/Mykrobe-tools/mykrobe, and the parser script at https://github.com/ShigellaGenomics/mykrobeshig (details in **Methods**).

**Figure 2:**
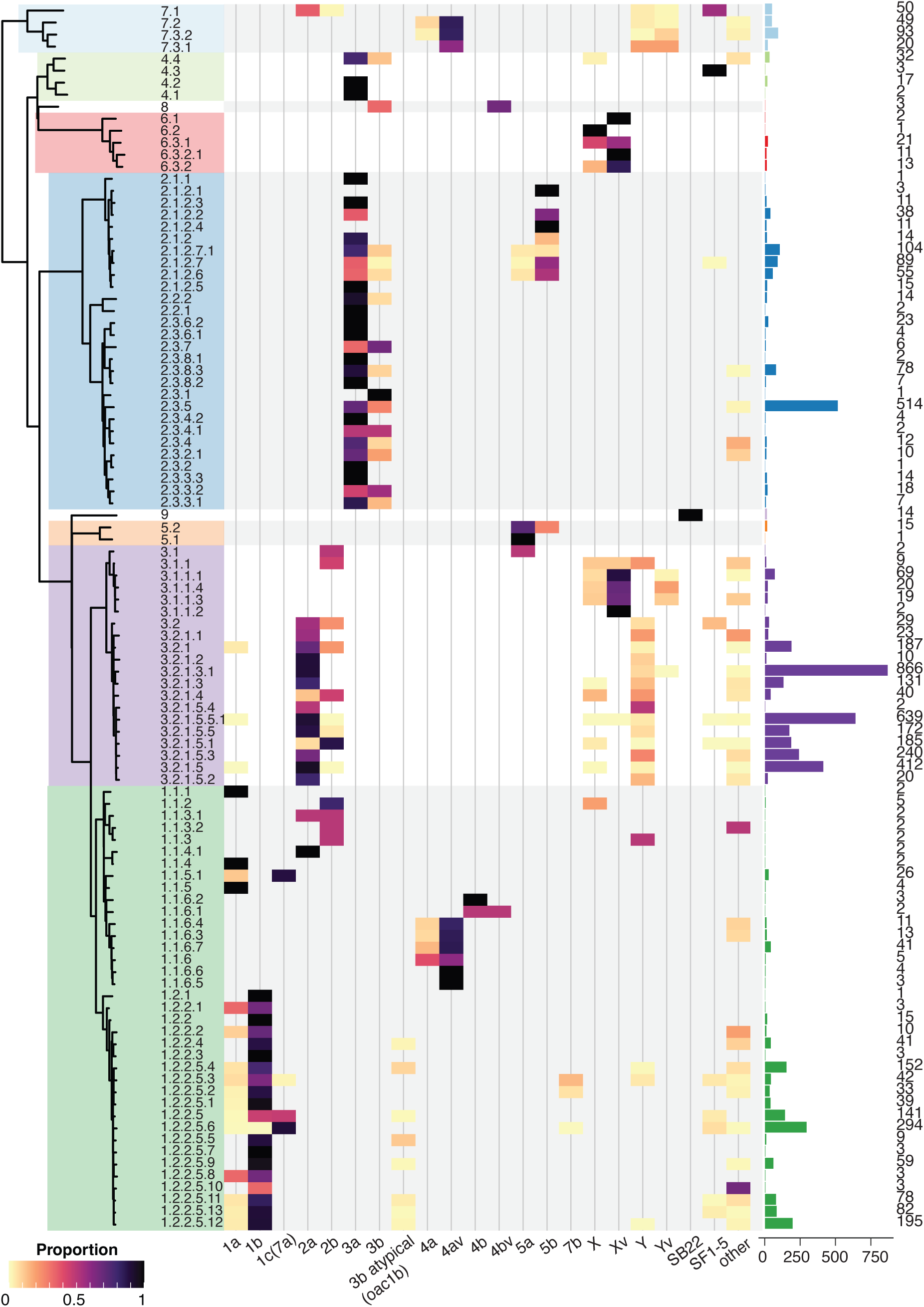
Genotyping scheme and correlation with serotype. Left, maximum-likelihood phylogeny of one representative per genotype. Shaded boxes behind phylogeny indicate PGs, with genotypes labelled next to the tree. Middle, heatmap showing the percentage of all retrospective genomes (discovery + validation datasets) that carry a particular serotype as predicted by ShigaPass. Right, bar plot indicates total number of genomes per genotype, with totals to the right of the bar, and bars coloured by PG as per phylogeny.

When we applied Flex-It to the full 2,000-genome ‘discovery dataset’ on which it was initially developed, we found Mykrobe demonstrated correct assignments of genotypes for 99.75% (n=1,994/2,000) of the isolates, most with either ‘strong’ (n=1,984) or ‘moderate’ (n=6) confidence. Nine genomes had ‘weak’ confidence for the calls, indicating that Mykrobe was unable to obtain confident support for one or more levels of the genotyping hierarchy. Of the nine ‘weak’ confidence genomes, five genomes were incorrectly genotyped by Mykrobe, however in all cases the correct genotype was being detected by Mykrobe, but at similarly low confidence (**Supplementary Table 3**).

We then used Flex-It to analyse the remaining retrospective 3,820 genomes that formed the ‘validation set.’ Cross-referencing the resulting genotype calls with the full maximum-likelihood phylogeny demonstrated Flex-It calls were correctly made (defined as being monophyletic with others of the same genotype) for all 3,820 genomes. Nearly all genomes were genotyped with ‘strong’ confidence (99.6%, n=3,808/3,820), and four genomes with ‘moderate’ confidence. The remaining eight genomes had ‘weak’ confidence calls, however all were genotyped correctly by Mykrobe, after inspecting the phylogeny (**Supplementary Table 3**).

To determine the utility of Flex-It for detecting point mutations in the QRDRs in GyrA-83, GyrA-87 and ParC-80, we calculated the concordance between our initial mapping-based approach (see **Methods**) and Flex-It Mykrobe calls for all 5,819 genomes in the curated dataset. Mykrobe correctly identified the same QRDR mutations as the mapping approach in 100% of cases, however, in six genomes Mykrobe called additional mutations in GyrA or ParC which our mapping approach did not detect.

As the Flex-It implementation in Mykrobe works with either short-read or long-read datasets, we validated the novel scheme against 18 publicly available genomes with matched Illumina and Oxford Nanopore Technologies (ONT) data (**Supplementary Table 4**). The ONT data included both legacy R9.4.1 chemistry (n=11), and the most up-to-date chemistry, R10.4.1 (n=7). Flex-It gave identical results in all 18 genomes, correctly identifying genotype and QRDR mutations (**Supplementary Table 4**), demonstrating the applicability of the scheme to different read-sets.

Finally, to test specificity, we tested the scheme and parser against 18 short-read sequenced genomes that belonged to different *Shigella* serogroups (see **Methods**), to check that the species probes and scheme were not cross-reactive against other *Shigella*. Our scheme correctly identified that these genomes belonged to different *Shigella* serogroups in all cases, flagging them as ‘Unknown *E. coli* / *Shigella*’ (**Supplementary Table 5**).

### Distribution of serotypes across the *S. flexneri* population

We explored the diversity of serotypes within each individual genotype. Thirty-four (of the 119) genotypes had only a single serotype (**Figure 2**, **Supplementary Table 1**), although only nine of these genotypes had ≥10 genomes. The trend of multi-serotype diversity in PGs (**Figure 1d**) continued at the individual genotype level, particularly in the three largest PGs 1, 2 and 3 (**Figure 2**). In PGs 1 and 3, there were substantial differences based on lineage. For example, subserotype 1b was detected at high frequency in all but three genotypes in lineage 1.2 (33-100% of genomes, except genotype 1.2.2.5.6), with other subserotypes (e.g. 1a, 3b atypical, 7b, Y and ‘*other’*) found at much lower frequencies (<30%), suggestive of continued recombination of surface antigens, or movement of serotype converting phages and plasmids within genotypes. In contrast, genotypes within lineage 1.1 were spread between subserotypes in serotype 4 (clade 1.1.16), serotype 1 (clade 1.1.1), and serotype 2 (clades 1.1.2 and 1.1.3) (**Figure 2**). In PG3, lineage 3.2 was associated with subserotype 2a. In contrast, lineage 3.1 was primarily associated with subserotype Xv or 2b (**Figure 2)**, which were only otherwise detected in PG7 (for subserotype 4a or 4av) or PG8 (for subserotype 4bv). PG2 was the outlier, where all but two genotypes had a high frequency of either subserotypes 3a or 3b, except for genotypes 2.1.2.1 and 2.1.2.4, where subserotype 5b predominated (**Figure 2**). In the genotypes within PGs 4 to 9, only five genotypes were associated with a single subserotype.

These were 4.1 and 4.2 (both associated with 3a, n = 2 and n = 17 genomes respectively), 5.1 (associated with 5a, n = 1), 6.1 (associated with Xv, n = 2), and 9 (associated with SB22, n = 13). The remaining 11 genotypes, though low in frequency, were characterised with two or more serotypes.

### AMR to therapeutic drugs has arisen in multiple genotypes

While the distribution of AMR varied within and between the different genotypes of *S. flexneri,* nearly all *S. flexneri* had at least one known AMR mechanism (94.7%, n = =5,511/5,819). The 308 genomes with zero AMR determinants belonged to one of nine genotypes, each of which had <5 genomes (in PGs 1, 2 and 3, **Figure 3**). The *Shigella* resistance locus (SRL, carrying AMR mechanisms against ampicillin (*bla*_OXA-1_), streptomycin (*aadA1*), chloramphenicol (*catA1*) and tetracycline (*tet(B*)) was prevalent across the diversity of *S. flexneri*, with 80/119 (67%) of genotypes carrying at least one SRL AMR gene, and 53/119 (44%) of genotypes with all four SRL AMR genes (**Figure 3**).

**Figure 3:**
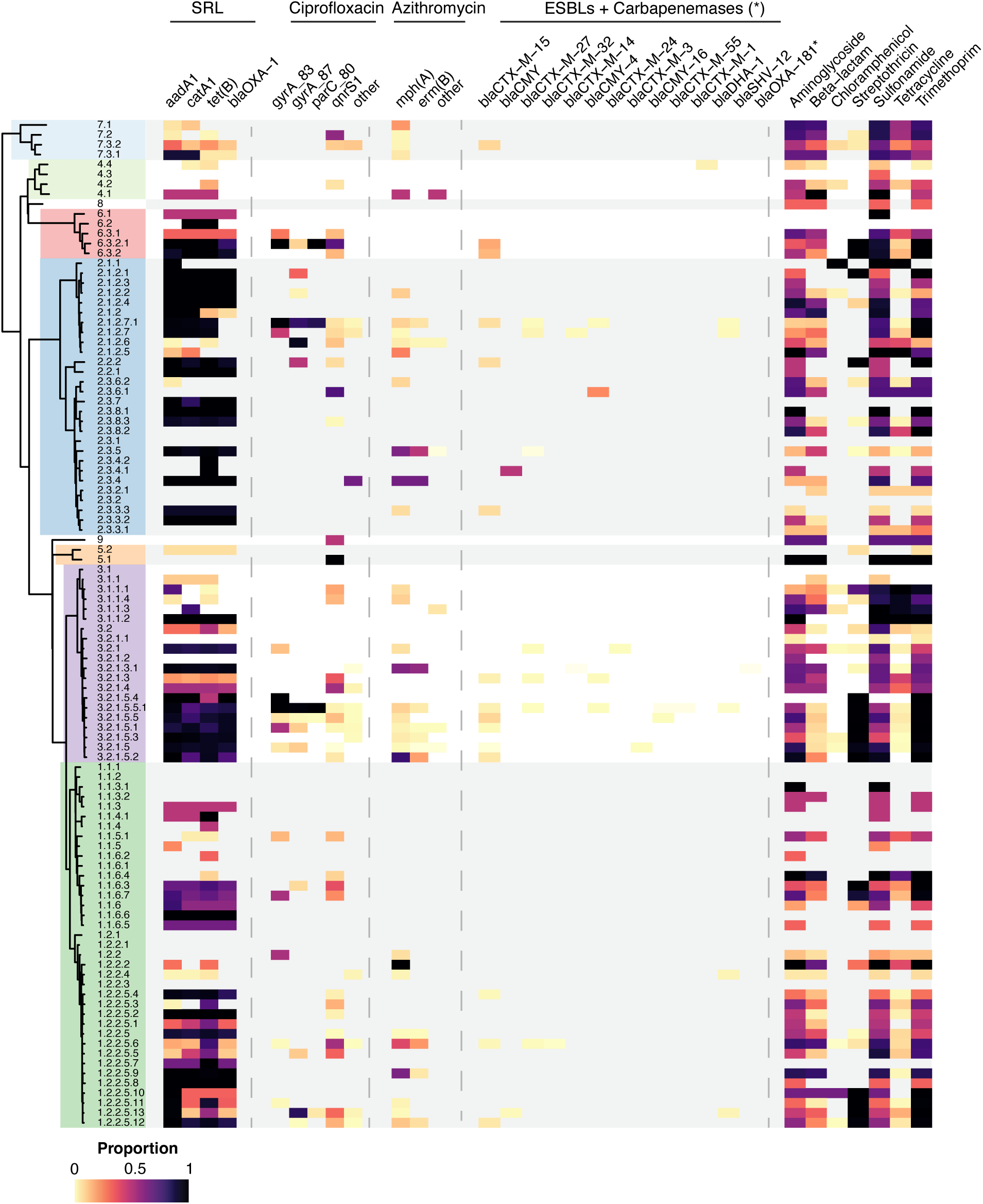
**AMR profiles associated with genotypes in 5,819 *Shigella flexneri*** Genotypes labelled and coloured by PG as per Figure 2. Heatmap shows the percentage of all genomes (discovery + validation) that carry a particular AMR marker, as per legend. *Shigella* resistance locus (SRL), ciprofloxacin (fluoroquinolones), azithromycin (macrolides) and ceftriaxone (3GCs) + carbapenemase determinants are shown individually, with the single carbapenemase gene, *bla*_OXA-181_, indicated by an asterisk (*). All other determinants are grouped by drug class and shown to the far right of the heatmap.

Remarkably, only three genotypes were associated with multiple point mutations in the QRDRs at high frequency (**Figure 3**) and all three were globally distributed, indicating the convergent evolution and spread of against first line fluoroquinolone treatment. Two genotypes had triple QRDR profiles, associated with clinical resistance to ciprofloxacin, at high frequency. These were genotype 2.1.2.7.1 (n = 104, with four different serotypes, **Figure 2**), which had 82% of genomes with GyrA-S83L + GyrA-D87G + ParC-S80I mutations together. The remaining 18% of genomes in 2.1.2.7.1 were either double QRDR mutants with GyrA-S83L + ParC-S80I, or single QRDR mutants with GyrA-S83L (**Figure 3**). Genotype 2.1.2.7.1 spanned multiple geographical regions in Europe, Asia and Oceania, with the earliest genomes dated from 2008. Of these eight 2008 genomes, seven were triple mutants, and one was a single GyrA-S83L mutant. Genotype 3.2.1.5.5.1 (n = 637, with >six different serotypes, **Fig 2**) had 98% of genomes with triple QRDR mutations in GyrA-S83L + GyrA-D87G/N + ParC-S80I, and 1.8% were double QRDR mutants with GyrA-S83L + ParC-S80I. Members of genotype 3.2.1.5.5.1 are globally disseminated, spanning two decades, with the earliest genome from 1999, which was a triple QRDR mutant.

This genotype has previously been identified as BAPS2 in an Australian study that included all *S. sonnei and S. flexneri* received at the state of Victoria reference laboratory^13^, but otherwise has not been identified as a major lineage in previous studies. The only double mutation genotype was 6.3.2.1 (n = 11, serotype Xv), which had GyrA-S83L and ParC-S80I mutations. These genomes and the isolates were collected from multiple geographical regions including UK, France, Australia and India and spanned a decade from 2011-2019 (**Supplementary Table 1**). These findings demonstrate the emergence of resistance to ciprofloxacin in multiple genotypes, which also have different serotypes within the genotype which impacts surveillance efforts.

AMR determinants for second line treatments azithromycin and ceftriaxone were detected in multiple genotypes (n = 34 and n = 24, respectively) at low frequency (median 6.4% and 1.12% of genomes within a genotype, respectively). However, four genotypes with n>15 genomes had the azithromycin resistance determinant, *mph(A),* in >50% of genomes (genotype 1.2.2.5.9, n = 59, 64%; genotype 2.3.5, n = 514, 64%; genotype 3.2.1.3.1, n = 863, 63%; genotype 3.2.1.5.2, n = 20, 80%). Three of these genotypes were associated with previously reported lineages, 1.2.2.5.9 (BAPS1 in Ingle *et al* 2019 ^13^), 2.3.5 (sublineage A/B/C and BAPS3 in Baker *et al* 2018 and Ingle *et al* 2019 ^13,14^), and 3.2.1.3.1 (BAPS2 in Ingle *et al* 2019^13^). Genotypes 2.3.5 and 3.2.1.3.1 were globally disseminated, while 1.2.2.5.9 was more associated with high income regions, and 3.2.1.5.2 was identified in isolates from Europe and North America.

Genes conferring ceftriaxone resistance were found in only 2% (n = 120/5,819) of genomes. Thirteen unique ceftriaxone resistance genes were detected, with *bla*_CTX-M-15_ (n = 82) and *bla*_CTX-M-27_ (n = 12) the most common (**Figure 3**). Of the 24 genotypes that harboured a ceftriaxone resistance gene, only five of these had ≥20 genomes total with ≥5% of these carrying a ceftriaxone resistance gene. Genotype 3.2.1.5.5, the parent to the triple QRDR mutation-containing genotype 3.2.1.5.5.1, contained genomes with both ceftriaxone and QRDR profiles from as early as 2011, in Pakistan^1,34^. The two determinants detected were *bla*_CTX-M-15_ (n = 15) and *bla*_CTX-M16_ (n = 1). Two genotypes with genomes carrying *bla*_CTX-M-15_ were 7.3.2 (n = 8/93, 8.6%) and 1.2.2.5.12 (n = 10/195, 5.1%). The 7.3.2 isolates were all collected in France and were serotyped as either 4av or Yv, with the earliest isolate from 2017 (**Supplementary Table 1**). The 1.2.2.5.12 isolates nearly all originated from Europe with the earliest isolate from 2016. These data demonstrate that acquired AMR mechanisms to important therapeutic drug classes have arisen in multiple genotypes.

### Flex-It provides high level resolution for public health surveillance

To explore the public health utility of Flex-It in analysing routinely generated *S. flexneri* data more broadly, we applied the novel scheme to 13,273 publicly available *S. flexneri* genomes from public health surveillance laboratories collected over a decade from 2015 to 2024 in three HICs; the UK, the USA and Australia. A total of 12,765 genomes (97%) were confidently genotyped with Flex-It. Of those that were not confidently called, eleven were removed at the serogroup’-call stage as ‘not *E coli/Shigella,* and a further 478 were not found to be *S. flexneri* so were called as ‘Unknown *E. coli/Shigella*’. Fifteen genomes were genotyped with ‘weak’ confidence, and four had no genotype detected at all, and were excluded from subsequent analysis.

Of the 119 genotypes in Flex-It, 83 were detected in the public data, demonstrating a continued high level of co-circulating diversity of genotypes across the three locations (**Supplementary Table 6**). Whilst each region has substantially different sampling and sequencing approaches for *S. flexneri*, we were able to identify common genotypes across the three regions (**Supplementary Figures 3-4, Supplementary Table 6**), demonstrating the utility of having a scheme with standardised nomenclature. We found that in PGs 1, 2 and 3, while the top genotypes were similar across all three regions (e.g. genotypes 1.2.2.5.6, 2.3.5 and 3.2.1.5.5.1, **Table 2, Supplementary Figures 3-4**), each region had a high diversity of genotypes detected, particularly the USA (Simpson’s Diversity Index 0.894 for USA vs 0.874 and 0.836 in the UK and Australia, respectively).

**Table 2:** Summary of diversity of genotypes detected in nine phylogroups in public data from three HICs over a ten-year period (2015-2024).

| Phylogroup | Country | Genomes (n) | Unique genotypes (n) | Simpson Index | Top 3 genotypes |
| --- | --- | --- | --- | --- | --- |
| 1 | Australia | 191/1204 | 16 | 0.78 | 1.2.2.5.6 (n=78)<br>1.2.2.5.12 (n=33)<br>1.2.2.5.13 (n=24) |
| 1 | USA | 2150/6105 | 24 | 0.75 | 1.2.2.5.6 (n=905)<br>1.2.2.5.12 (n=382)<br>1.2.2.2 (n=375) |
| 1 | UK | 1836/5456 | 30 | 0.68 | 1.2.2.5.12 (n=939)<br>1.2.2.5.6 (n=398)<br>1.2.2.5.4 (n=83) |
| 2 | Australia | 137/1204 | 10 | 0.69 | 2.3.5 (n=70)<br>2.1.2.7.1 (n=18)<br>2.3.4 (n=18) |
| 2 | USA | 1281/6105 | 10 | 0.58 | 2.3.5 (n=594)<br>2.3.4 (n=578)<br>2.1.2.7.1(n=83) |
| 2 | UK | 860/5456 | 16 | 0.49 | 2.3.5 (n=600)<br>2.1.2.7.1(n=106)<br>2.3.8.3 (n=53) |
| 3 | Australia | 835/1204 | 12 | 0.68 | 3.2.1.5.1 (n=406)<br>3.2.1.5.5.1 (n=185)<br>3.2.1.3.1 (n=147) |
| 3 | USA | 2652/6105 | 15 | 0.70 | 3.2.1.5.5.1 (n=1306)<br>3.2.1.5.2 (n=457)<br>3.2.1.3.1 (n=297) |
| 3 | UK | 2643/5456 | 19 | 0.67 | 3.2.1.5.5.1 (n=1283)<br>3.2.1.3.1 (n=772)<br>3.2.1.3 (n=152) |
| 4 | Australia | 7/1204 | 3 | 0.44 | 4.4 (n=5)<br>4.1 (n=1)<br>4.2 (n=1) |
| 4 | USA | 7/6105 | 1 | 0.00 | 4.4 (n=7) |
| 4 | UK | 16/5456 | 3 | 0.40 | 4.4 (n=12)<br>4.3 (n=3)<br>4.2 (n=1) |
| 5 | UK | 3/5456 | 1 | 0.00 | 5.2 (n=3) |
| 6 | Australia | 6/1204 | 1 | 0.00 | 6.3.2.1 (n=6) |
| 6 | UK | 16/5456 | 2 | 0.49 | 6.3.2.1 (n=9)<br>6.3.2 (n=7) |
| 7 | Australia | 27/1204 | 4 | 0.63 | 7.2 (n=14)<br>7.3.1 (n=6)<br>7.1 (n=6) |
| 7 | USA | 14/6105 | 2 | 0.34 | 7.3.2 (n=11)<br>7.2 (n=3) |
| 7 | UK | 59/5456 | 4 | 0.53 | 7.2 (n=34)<br>7.3.2 (n=22)<br>7.3.1 (n=2) |
| 8 | UK | 2/5456 | 1 | 0.00 | 8 (n=2) |
| 9 | Australia | 1/1204 | 1 | 0.00 | 9 (n=1) |
| 9 | USA | 1/6105 | 1 | 0.00 | 9 (n=1) |
| 9 | UK | 21/5456 | 1 | 0.00 | 9 (n=21) |

We next investigated the application of Flex-It to public health data in combination with detection of additional AMR mechanisms using AMRFinderPlus (see **Methods**) to the key therapeutic agents ciprofloxacin, azithromycin and ceftriaxone, that confer the triple resistance profile associated with XDR *Shigella* (**Figure 4**). The proportion of genomes with predicted resistance ranged between 7-43%, depending on drug (**Figure 4a**). Genotype 3.2.1.5.5.1 was most commonly associated with ciprofloxacin resistance across all three regions (**Figure 4b**).

**Figure 4:**
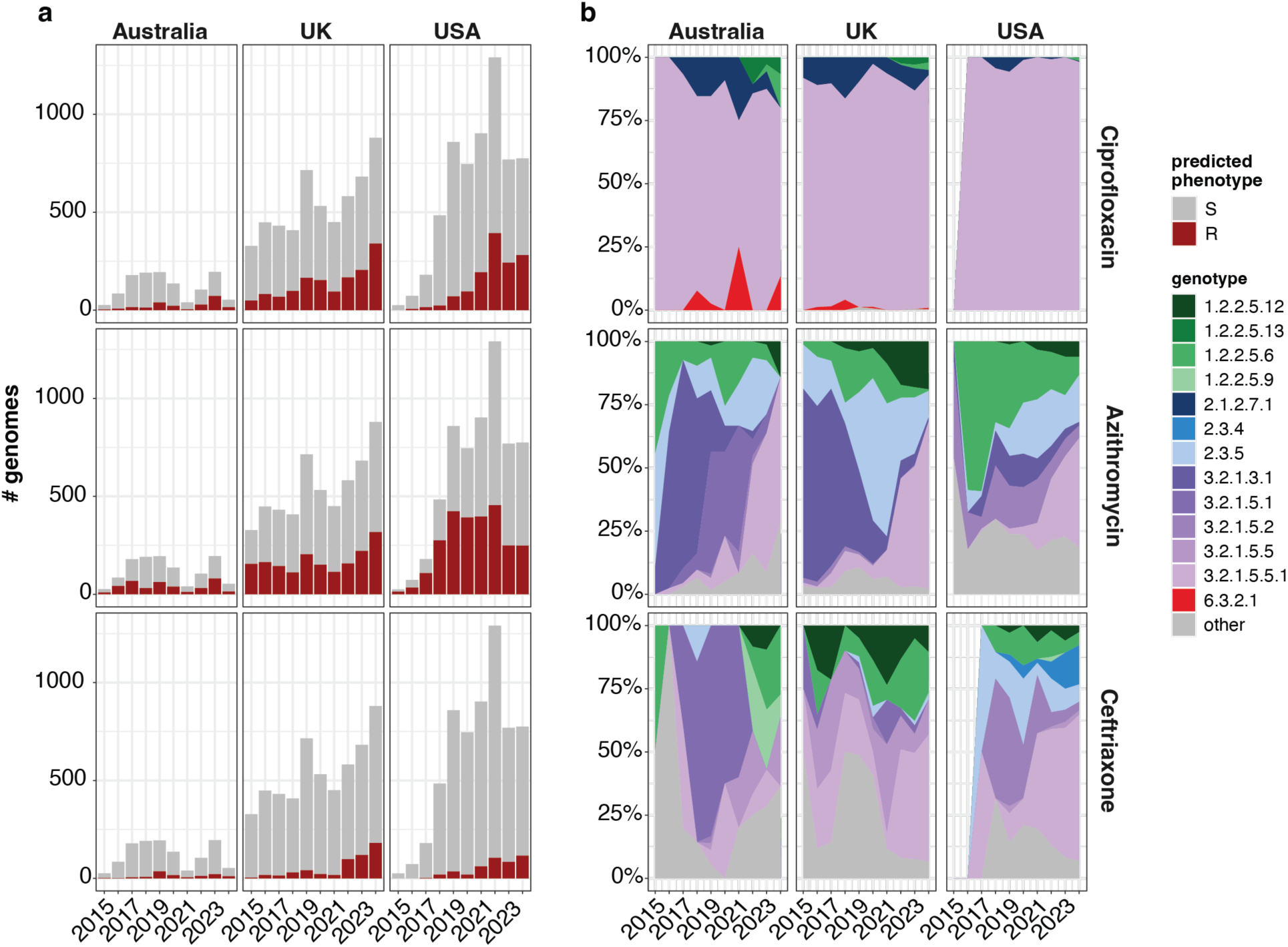
AMR profiles for empiric therapeutics and genotypes of genomes from three HICs. **a**) Barplots of the number of genomes with predicted phenotype resistance profile to three drug classes, ciprofloxacin (fluoroquinolones), azithromycin (macrolides) and ceftriaxone (3GCs), for the three HICs. **b**) Area plot showing the percentage of genomes by genotype, per year, across the three HICs for the three drug classes, ciprofloxacin, azithromycin and 3GCs. In the absence of any genomes identified as resistant, no genotypes are shown on the plot.

Genomes with predicted resistance to azithromycin and ceftriaxone came from a wider range of genotypes, however the most common genotypes were from the epidemiologically dominant PGs 1, 2 and 3, as expected based on recent outbreaks reported in HICs being driven by genotypes within these PGs^15,24,41,45^ . The four genotypes identified in the retrospective dataset carrying *mph(A)* genes, 1.2.2.5.6, 2.3.5, 3.2.1.3.1 and 3.2.1.5.2, were also detected in this public health dataset across all three HICs. Interestingly, genotype 3.2.1.5.5.1 was found to be harbouring AMR determinants towards azithromycin and ceftriaxone, in addition to ciprofloxacin in all three regions (**Figure 4b**).

We detected 330 XDR genomes (2.6%), with resistance mechanisms to all three agents in six different genotypes. Most of these genomes were at low frequency in their respective genotypes (1.2.2.5.13 (n=1), 1.2.2.5.6 (n=6), 2.1.2.7.1 (n=6), 3.2.1.5.5 (n=2), and 6.3.2.1 (n=4)), however, the majority of XDR genomes were found in 3.2.1.5.5.1 (n=314). Given that this genotype can easily obtain plasmids carrying azithromycin or ceftriaxone resistance determinants is concerning, and XDR outbreaks have now been reported during and prospective to our sampling frame^17^. Overall, while the data we have collated here isn’t suited to detailed genomic epidemiological analysis of the emergence or prevalence of AMR across the three regions, our data did indicate a likely increase in AMR across the sampling frame and the utility of having more precise nomenclature for important genotypes circulating.

## Discussion

We have developed an open-source, novel genotyping method for *S. flexneri*: Flex-It. In doing so, we have gained several insights into the structure and diversity of the globally circulating population of serogroup *S. flexneri* 1-5, X & Y by combining our scheme with geographical, temporal and accessory genome data such as AMR and serotype. We applied Flex-It to a curated retrospective dataset of 5,819 *S. flexneri* genomes, and to >13,000 publicly available genomes from public health laboratories from three HICs, demonstrating diversity of the 119 newly defined genotypes across time and geography. Moreover, these historical and comprehensively characterised *S. flexneri* genomes, and associated underlying data, will be an important resource to future studies where well-curated data are required for method and tool development.

We demonstrated that while serotyping was useful in combination with genotypes, grouping by serotype alone for epidemiological investigations can create noise in the data by including genomically unrelated organisms. We showed the diversity of predicted serotypes at both PG and individual genotype-level in the retrospective *S. flexneri* dataset, bringing further granularity to the previously described genomic variability and highlighting the high frequency of potential serotype switching in *S. flexneri*^34^. These insights provide a basis for future detailed investigation of serotype switching within genotypes of concern. The importance of moving away from serotype-driven outbreak investigation is further exemplified in recent studies that sought to track the emergence of MDR/XDR *S. flexneri* in different populations^20,23,24^. For example, in a 2023 study exploring the re-emergence of MDR *S. flexneri* 3a, the lineage of concern (genotyped as 2.3.5), was responsible for an increase in infections in the UK^24^.

However other genotypes, including those in different PGs, were also included in *S. flexneri* 3a genomes, highlighting that detecting this outbreak would have been more readily facilitated by the application of a genotyping scheme with enhanced genomic resolution^24^. In this increasingly common scenario, when a new AMR profile of concern emerges, Flex-It will provide a standardised framework based on genetic relatedness that will become foundational to public health genomic surveillance efforts^17^. Moreover, despite the global burden of shigellosis, there is no licensed vaccine for *Shigella,* with vaccine candidates in development typically O-antigen based^33,46–48^, an approach known to lead to strain replacement in *Streptococcus pneumoniae*^49,50^. Linking genotypes of concern and antigenic diversity will be important for current and future surveillance efforts to support vaccine development and implementation, and other novel therapeutics.

Ciprofloxacin-resistant *Shigella* is currently listed as WHO priority pathogen. Triple QRDR mutations in *S. flexneri* differ from what has been observed in *S. sonnei.* In *S. sonnei,* mutations leading to ciprofloxacin resistance arose in one parent genotype, with all descendant genotypes maintaining the triple-point QRDR profile^40^. In contrast, triple point QRDR mutation profiles have arisen in several *S. flexneri* genotypes, notably 2.1.2.7.1 and 3.2.1.5.5.1, with single point mutations widespread across the population. This potentially reflects the broader phylogenetic space occupied by *S. flexneri*, where the AMR evolution within a single PG may be more comparable to *S. sonnei*, and may reflect differences in the ecologies between the two species. The pattern observed in *S. flexneri* here is more akin to what is observed in *S.* Typhi where point mutations in QRDR associated with ciprofloxacin non-susceptibility are found in most genotypes^38,51,52^.

Additionally, we demonstrate that AMR mechanisms to azithromycin and ceftriaxone are highly prevalent and widely distributed within *S. flexneri.* Importantly, we observed that AMR genes predicting resistance to all key treatments are emerging convergently (i.e. in multiple genotypes) across the population structure. It is likely that *S. flexneri* will continue to acquire AMR genes, particularly those that are shown to be plasmid mediated and drive outbreaks in different settings as the horizontal acquisition of resistance among *Shigella* sp. has been repeatedly demonstrated^18,20,21,31,36,53^. Hence, a shared nomenclature that enables clear communication about the temporal trends of genotypes circulating within and between different international borders will be critical in tracking what lineages are circulating and what AMR profiles are present.

Multiple genotypes of concern identified in the retrospective dataset were also present in the public health dataset. The application of Flex-It to routinely collected public health data demonstrates the use of genotyping for both detecting lineages of concern and potentially quantify the impact of interventions. For example, the decrease in the number of *S. flexneri* genomes and genotype diversity in Australia between 2020 and 2022 corresponds to the public health measures introduced in Australia in this period (social distancing, lockdowns, closure of international and state borders) aimed at controlling COVID-19. This has been shown to impact other circulating bacterial pathogens^54,55^, and demonstrates how Flex-It revealed replacement of genotypes circulating in Australia before and after these measures. This will be key for quantifing changes in the population in response to other interventions such as potential vaccine rollouts, similar to those currently being undertaken for *S.* Typhi^56^.

A limitation of this work is that a more detailed genomic epidemiological investigation into the genotypes and AMR mechanisms to empiric therapies circulating in the three HICs was beyond the scope of this study. While these data were suited to demonstrate the broad applicability of Flex-It, the lack of detailed sampling and selection for WGS information from each of the three countries means linking genotype and AMR profiles, and exploring global circulation, will be a future avenue of research, now readily facilitated by our scheme. This genomic framework will provide the community with a standardised approach to monitor the spread of known *S. flexneri* drug-resistant genotypes, identify population expansions that may be driven by novel AMR profiles in existing genotypes and provide the tools to facilitate communication for public health surveillance without requiring the sharing of underlying genomic data.

This study has several strengths. First, it provides a robust and consistent nomenclature for an important priority pathogen that is publicly available and readily implementable. Flex-It is implemented in the same tool as the *S. sonnei* scheme, enabling a single workflow that covers >90% of shigellosis cases in most settings^57,58^. The Flex-It framework is amendable to adding new genotypes. For example, a new genotype(s) could be added should carbapenem resistance arise and expand in different genomic backgrounds of *S. flexneri,* which is anticipated to drive future outbreaks. The addition of new genotypes has been successfully undertaken in other genotyping schemes such as *S.* Typhi^38,59,60^. New genotype addition to Flex-It is planned to replicate efforts currently being undertaken for *S.* Typhi, by developing standardised criteria for new genotypes, including epidemiological and geographical considerations, and the continued use of consistent methods for defining marker SNPs.

Finally, a limitation of Flex-It is that it is only able to genotype *S. flexneri* serotypes 1-5, X & Y genomes. It cannot genotype *S. flexneri* serotype 6, which is not phylogenetically related to *S. flexneri* serotypes 1-5, X & Y genomes (CC245), but is still currently considered *S. flexneri.* Serotype 6 represents a different major lineage that emerged at a different time in the evolution of *E. coli,* and was identified as *S. flexneri* before the advent of WGS^5,19^. This polyphyletic quirk of *Shigella* nomenclature, and its impact on developing SNP based genotyping schemes is not limited to *S. flexneri* and future planned efforts to develop schemes for *S. boydii* and *S. dysenteriae* will also face this limitation, as these serogroups have several independent lineages that have emerged from an *E. coli* background. However, given that >90% of *S. flexneri* genomes analysed in the retrospective dataset were not serotype 6, and these are associated with large outbreaks and changing infection dynamics, Flex-It represents a significant advance in genomic surveillance of *Shigella*.

### Conclusions

Flex-It provides a novel and scalable approach for enhanced genomic surveillance of an important enteric pathogen, where ciprofloxacin-resistant *Shigella* are now listed as a WHO AMR priority pathogen. Similar schemes developed for other enteric pathogens, including *S. sonnei, Salmonella* Typhi and *Salmonella* Paratyphi B, have proven to be an important tool in tackling this public health threat posed by these drug-resistant enteric pathogens. Flex-It will address an identified gap by the *Shigella* research community in providing a genotyping scheme for *S. flexneri* serotypes 1–5, X and Y, informing public health surveillance and control strategies in an era of intense AMR emergence and vaccine development and implementation.

## Methods

### Selection of retrospective dataset and curation of metadata

We sought to capture the full diversity of *S. flexneri* circulating globally over several decades. We performed a literature search for studies containing *S. flexneri* genomes, published prior to 2023, where isolate metadata was provided. A full list of studies that were included in our dataset can be found in **Table 1**. Three genomes were removed because their read data was no longer available in the public archives; genomes that were present across multiple studies were de-duplicated.

Metadata for each genome was extracted from the supplementary files of each study where available, including year and country of isolation. If provided, we extracted information about whether the isolate was linked to an MSM outbreak. Reported travel information was used to determine if an isolate was collected from a high-income country but likely acquired from another region. If so, this was recorded as a travel associated isolate, and the country of isolation was updated to the likely country of origin. We additionally recorded any study-defined lineages for each isolate.

### Phylogenetic analysis

All genomes were mapped to the chromosome and plasmid of *S. flexneri* 2a str. 301 (accession AE005674.2) using RedDog v1.b11 (https://github.com/katholt/reddog-nf). Briefly, RedDog maps reads with Bowtie2 v2.2.9^61^ using the --sensitive-local parameter, then uses SAMtools v1.1^62^ to keep high quality SNP calls (phred score 20, read depth >5, heterozygous calls removed). Genomes passed mapping quality control if their read depth was >10x and >50% of reads in the sample mapped; 80 genomes were removed as they failed mapping. A further 14 genomes were excluded as they had a high ratio of heterozygous to homozygous SNPs (>0.7).

To generate the SNP alignment, SNPs called in repeat regions such as insertion sequences and phage were masked (**Supplementary Table 7**). Recombinant SNP sites were detected with Verticall v0.4.2 (https://github.com/rrwick/Verticall) using the alignment tree workflow. The final alignment consisted of 86,064 SNPs (**Supplementary File 1**). We used IQ-TREE v2 to infer a maximum likelihood phylogeny of all genomes using a GTR+G substitution model and 1000 bootstrap replicates^63^ (**Supplementary File 2**).

The phylogeny revealed 592 genomes that belonged to *S. flexneri* serotype 6, *S. boydii*, or *S. sonnei*, which were removed from further analysis. This left a final retrospective dataset of 5,819 genomes. A full list of all genomes included in the study, and their associated metadata, can be found in **Supplementary Table 1**. The full ML phylogeny with associated metadata has been uploaded to Microreact (https://microreact.org/project/sWtrZuAeQLQWZcNyQamX8a-shigella-flexneri-genotyping-scheme).

### Defining discovery and validation datasets

We split the retrospective dataset into two segments - an initial ‘discovery dataset’, for defining the genotyping scheme, and a ‘validation dataset’, for testing the genotyping scheme. The discovery dataset consisted of 2,000 genomes: 1,999 diverse genomes were selected with Treemmer v0.3^64^, plus the reference genome *S. flexneri* 2a str. 301. The validation dataset consisted of the remaining 3,820 genomes.

### Determining serotype and AMR profiles

The retrospective dataset of 5,819 genomes was assembled with Unicycler^65^ v0.5.1 using default parameters. We determined in-silico serotype from the genome assemblies with ShigaPass^42^ v1.5.0. A total of 203 genomes failed in-silico serotype detection, likely due to poor assembly quality. ShigaPass had been optimised for use with Spades assemblies, so all 203 genomes were re-assembled with Spades v4.2 using the parameters --careful --cov-cutoff auto -k 21,33,55,77 as per Yassine *et al*^41^. The Spades assemblies were then re-typed with ShigaPass, and serotypes were successfully predicted for 202/203 assemblies. AMR genes were detected from the original Unicycler assemblies using AMRFinderPlus^66^ v3.11.4 with database v2023-02-23.1 using default parameters. Mutations in the QRDRs of *gyrA* and *parC* were extracted from the read mapping-based SNP calls described above.

### Determining clades and subclades of the genotyping scheme

The discovery dataset was used to define the genotyping scheme. First, genomes were assigned to PGs based on Connor *et al*^19^, and their position within the phylogeny. We overlaid this with study-defined lineages where available. cgMLST profiles, where available, were extracted for all genomes using EnteroBase^37^. To define phylogroups, we aimed to retain backwards compatibility with the original PGs from Connor *et al.* We used HierCC clusters at the HC1100 level, as described in Yassine *et al*^41^, to define PGs, including the new PG9. The exceptions were PGs 1, 3, 4 and 8, where the HC1100 level was insufficient to split these into monophyletic groups. The HC400 threshold is sufficient to define PG3 separately from PG1. The original PG4 was not monophyletic, and therefore incompatible with a genotyping scheme, so we manually defined PG8 as the smaller branch of this group to retain monophyly across all PGs in the scheme. For PGs 4-9, HC400 levels were used to define lineages within these phylogroups. Due to the small number of genomes within these PGs, clades and subclades were defined by visual inspection of the phylogeny, using the epidemiological metadata and known study clusters.

For the large phylogroups PG1, PG2 and PG3, we used similar methods to Hawkey *et al* ^40^ to define clades and subclades within lineages. We calculated pairwise SNP distances within each PG using the C++ implementation of pairsnp v0.3.1 (https://github.com/gtonkinhill/pairsnp) from the recombination-free alignment. We inspected the pairwise SNP distributions and clustered dendrograms made from these distributions using single-linkage clustering with *hclust* in R, to identify sensible cutoffs that could be used to define lineages, clades and subclades (**Supplementary Figure 2**). Generally, we used troughs in the pairwise distributions to set cutoffs, with the exception of subclade in PG1, where there was no trough at the right location. Therefore, in this one instance, we used the dendrogram to guide cutoff selection (**Supplementary Figure 2**).

Genotypes were only defined if the cluster included at least two genomes. All genotypes were checked to ensure they were monophyletic, using the *getMRCA* function in ape^67^ v5.7-1 and the *Ancestors* function in phangorn^68^ v2.12.1. Any non-monophyletic clusters were manually resolved, which in some cases required subclades to be either merged, split into two groups, or needed additional groupings below the subclade level to enforce monophyletic relationships. To assign SNPs to genotypes for the scheme, we mapped the SNPs back onto the phylogeny using SNPpar^69^ v1.2 using default parameters (**Supplementary Files 3-5**). SNPs that were in core genes, with a non-synonymous:synonymous ratio of <1 were prioritised, as per Hawkey *et al*^40^. The final selection of SNPs and their locations and non-synonymous:synonymous ratio is in Supplementary Table 2.

### Implementation of genotyping scheme in Mykrobe

The genotyping scheme was implemented in Mykrobe^43,44^ v13.0, which uses a *k*-mer based approach to identify marker SNPs. This software has been utilised for genotyping in several enteric bacterial pathogens, including *S. sonnei*^40^, *S.* Typhi^38^ and *S.* Paratyphi B^39^, and previously benchmarked to run <1 min per genome. Mykrobe probes were created using the *mykrobe variants make-probes* command using a *k*-mer size of 21 and the *S. flexneri* 2a str. 301 reference genome (AE005674.2). This probe set also includes probes for the detection of mutations in the QRDRs of *gyrA* and *parC*. Mykrobe first determines whether a genome belongs to the *E. coli/Shigella* complex by looking for the presence of the *uidA* gene, and then if it is an *S. flexneri* genome by checking if it matches one of the 61 known STs for *S. flexneri*. We developed the *mykrobeshig* parsing script (available at https://github.com/ShigellaGenomics/mykrobeshig) to parse the Mykrobe output. The parser applies a confidence score to the genotype call: ‘strong’ indicates that Mykrobe has a high confidence in all levels of the genotype; ‘moderate’ indicates reduced confidence of 0.5 in only one level of the genotype; ‘weak’ indicates reduced confidence of 0.5 in two or more levels.

To ensure that an input genome belongs to the correct *S. flexneri* 1-5, X & Y clade, the parser script checks the ‘species coverage’ output by Mykrobe. If this value is >85.5%, then this genome belongs to the correct *S. flexneri* group. We set this threshold after running Mykrobe with the Flex-It scheme against 18 *Shigella* genomes from different serogroups (n = 4 for *S. sonnei* and *S. dysenteriae*, n = 5 for *S. boydii, S. flexneri* serotype 6 and *E. coli,* accessions in **Supplementary Table 5**). In all cases, matches to the *S. flexneri* species probes were <85.5% (**Supplementary Figure 5**). The full *S. flexneri* probe set can be found at https://doi.org/10.26180/31872106, and the code and input files for generating the probes are at https://github.com/ShigellaGenomics/flexit_probes.

Finally, Flex-It was run on 18 publicly available isolates with both long-read ONT data and short read Illumina data (accessions can be found in **Supplementary Table 4**). The ONT data included reads generated with either R9.4.1 or R10.4.1 chemistry. We compared the results of the Illumina genotype calls to the ONT genotype calls to determine concordance between sequencing technologies.

### Application to three unbiased datasets spanning 2015-2024

We collated a set of publicly available genomes, sequenced by three public health surveillance labs in Australia (Microbiology Diagnostic Unit, Victoria), the UK (UK Health Security Agency) and the USA (Centre for Disease Control) that do routine sequencing of *Shigella*. We selected data by accessing the NCBI Pathogen Detection Isolate Browser on August 29th, 2025, and filtered for genomes matching the following criteria: i) species taxid of 623 (*E. coli/Shigella*); ii) collection date between January 1st, 2015 and December 31st, 2024; iii) sequencing platform was Illumina; and iv) the collected_by lab was either CDC (for the USA), PHE (for UKHSA) or MDU-PHL (for Australia).

This gave us a set of 13,273 genomes which were typed using the Flex-It scheme in Mykrobe. We detected additional AMR determinants from assemblies, using AMRFinderPlus on assemblies generated with Unicycler, as described above for the retrospective dataset. We defined ciprofloxacin resistance as the presence of triple QRDR mutations, or double QRDR mutations plus a *qnrA* gene. Azithromycin resistance was defined as the presence of *mph(A)*, and ceftriaxone resistance as the presence of a 3GC gene. A full list of genomes, their accessions, genotype calls from Mykrobe, and resistance determinants to ciprofloxacin, azithromycin and ceftriaxone can be found in **Supplementary Table 6**.

### Data availability

All code and underlying data are made available through the supplementary material, GitHub, FigShare (https://doi.org/10.26180/31697419) and Microreact (https://microreact.org/project/sWtrZuAeQLQWZcNyQamX8a-shigella-flexneri-genotyping-scheme).

## Funding

JH was supported by an Emerging Leadership Fellowship from the National Health and Medical Research Council (NHMRC) of Australia (2034741). BPH is supported by a Leadership Fellowship from the NHMRC of Australia (2041625). DJI was supported by an Emerging Leadership Fellowship from the NHMRC of Australia (2041653) and the Research Accelerator Fund - Bioinformatics 2024 generously supported by Doherty Institute philanthropic donations.

## Acknowledgements

This work was supported by Monash eResearch capabilities, including the high-performance computer M3 and Research Data Storage.

## Notes

### Competing Interest Statement

The authors have declared no competing interest.

