## Supplementary Material for "Flex-It: A global standardised genotyping framework for *Shigella flexneri*"


### Supplementary Information

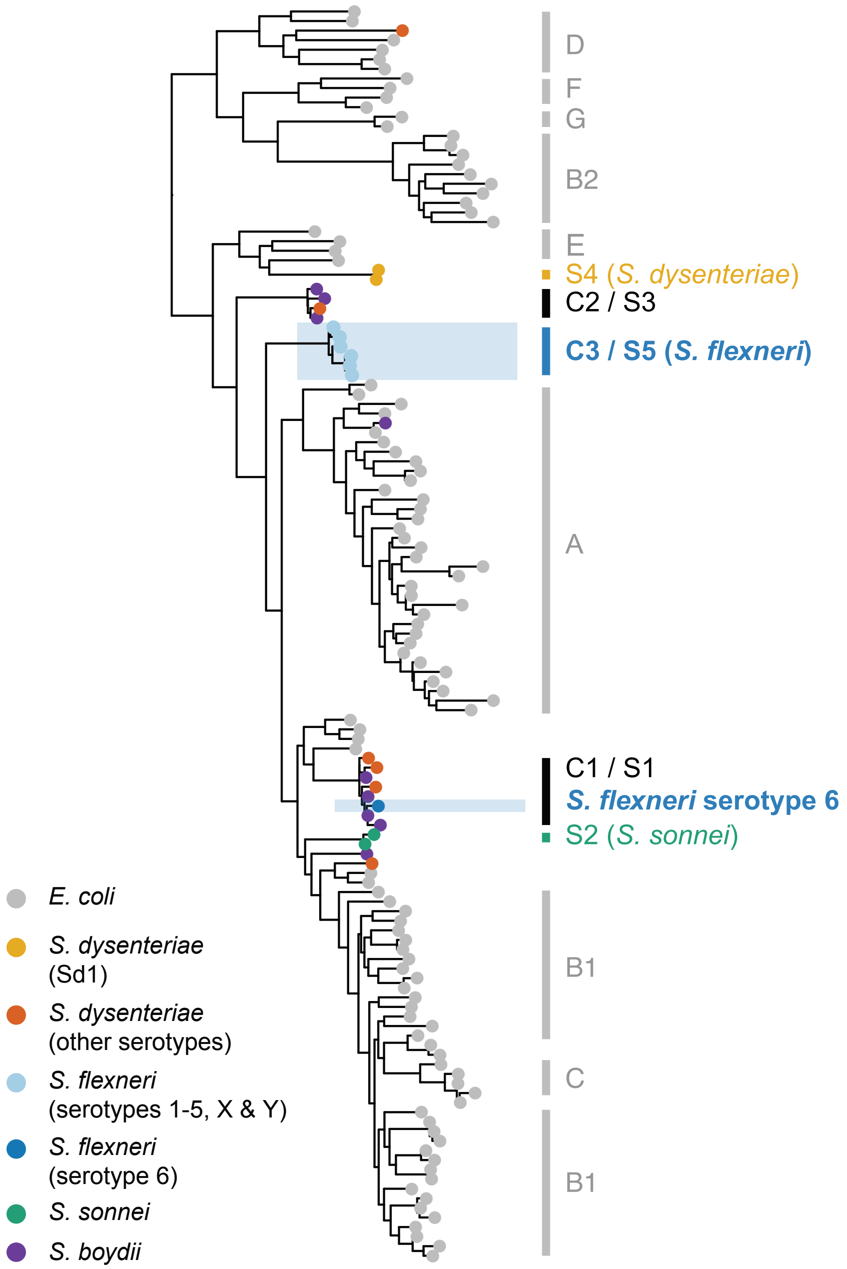


**Fig S1: Overview of *E. coli* and nested *Shigella* species**

Framework phylogeny of *E. coli* and nested *Shigella* serogroups (adapted from Hawkey *et al* 2020). The grey bars and letters to the right of the phylogeny refer to high level phylogroups within *E. coli* (as defined by Clermont *et al* 2013). Coloured tips of the tree indicate *Shigella* serogroups*.* The coloured blocks and letters to the right indicate previous nomenclature used across *Shigella.* For example, the C3 / S5 is the clonal complex 245 of *S. flexneri,* with members carrying the same chromosomal *rfb* gene cluster and various O-antigen converting prophages and plasmids. The distinct *S. flexneri* clade nested in C1/S1 with *S. boydii* and some serotypes of *S. dysenteriae* represents serotype 6 *S. flexneri* and is an independently evolved genomic lineage (clonal complex 145)

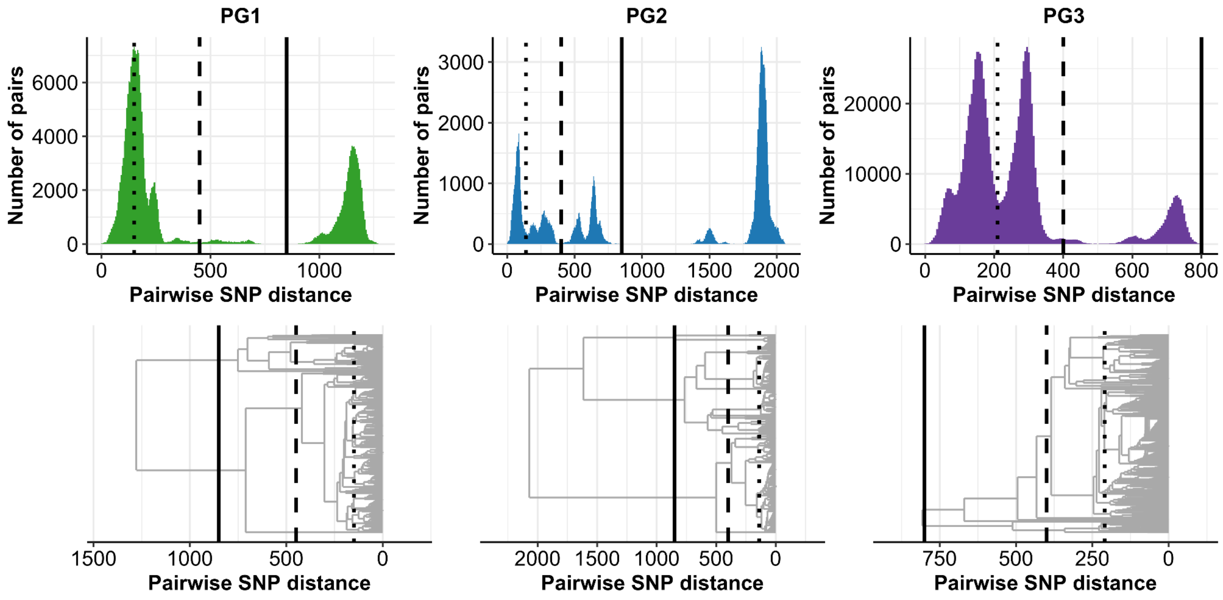


**Fig S2: Pairwise SNP distances for PGs 1, 2 and 3.**

Top, Distribution of pairwise SNP distances for each PG. Lines indicate cutoffs for lineage (solid line), clade (dashed line) and subclade (dotted line). Bottom, Clustered dendrograms of pairwise SNP distances for each PG, with cutoffs for lineage, clade and subclade indicated by black lines as per top row.

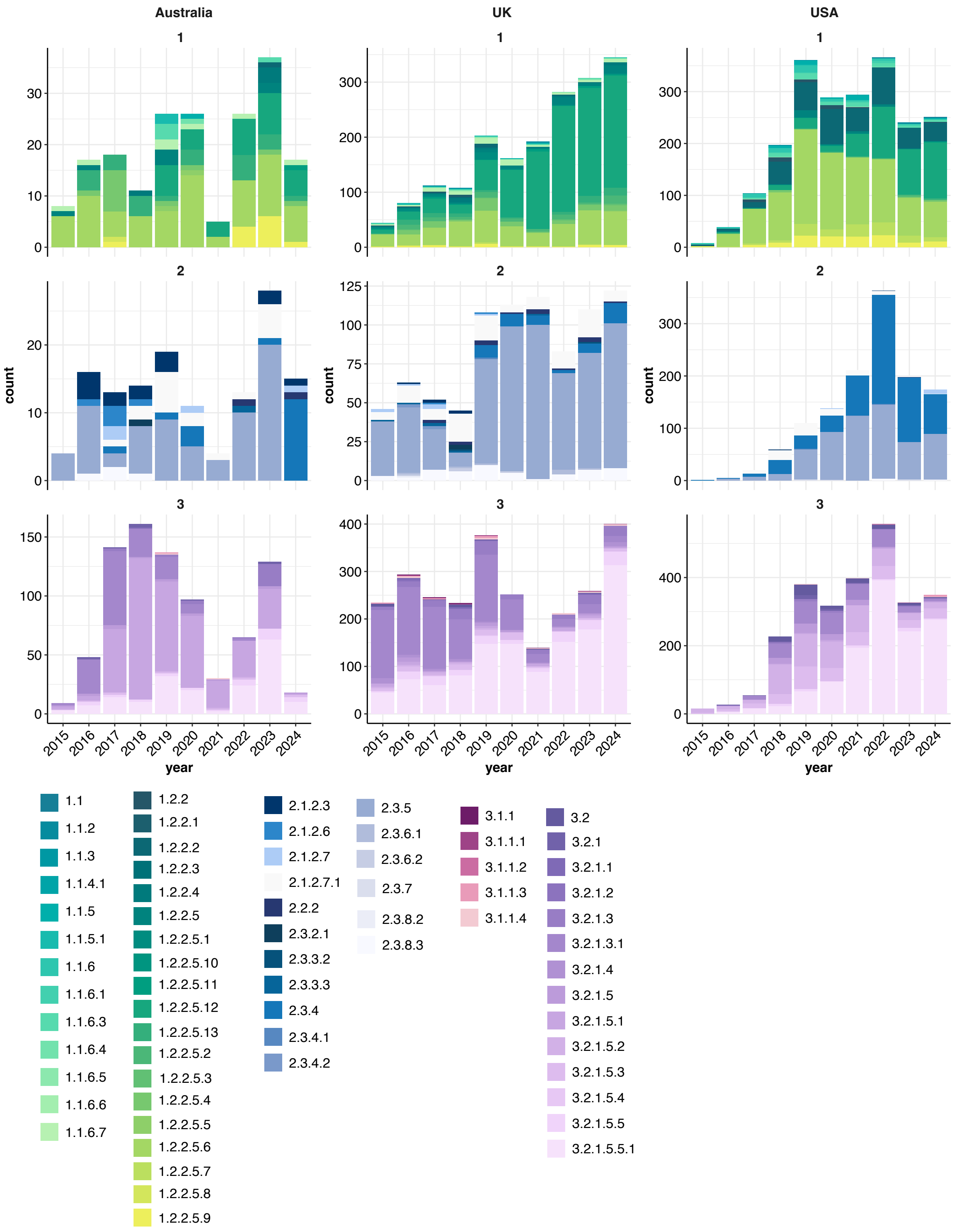


**Fig S3: Barplots of genotypes across three HICs for PGs 1-3**

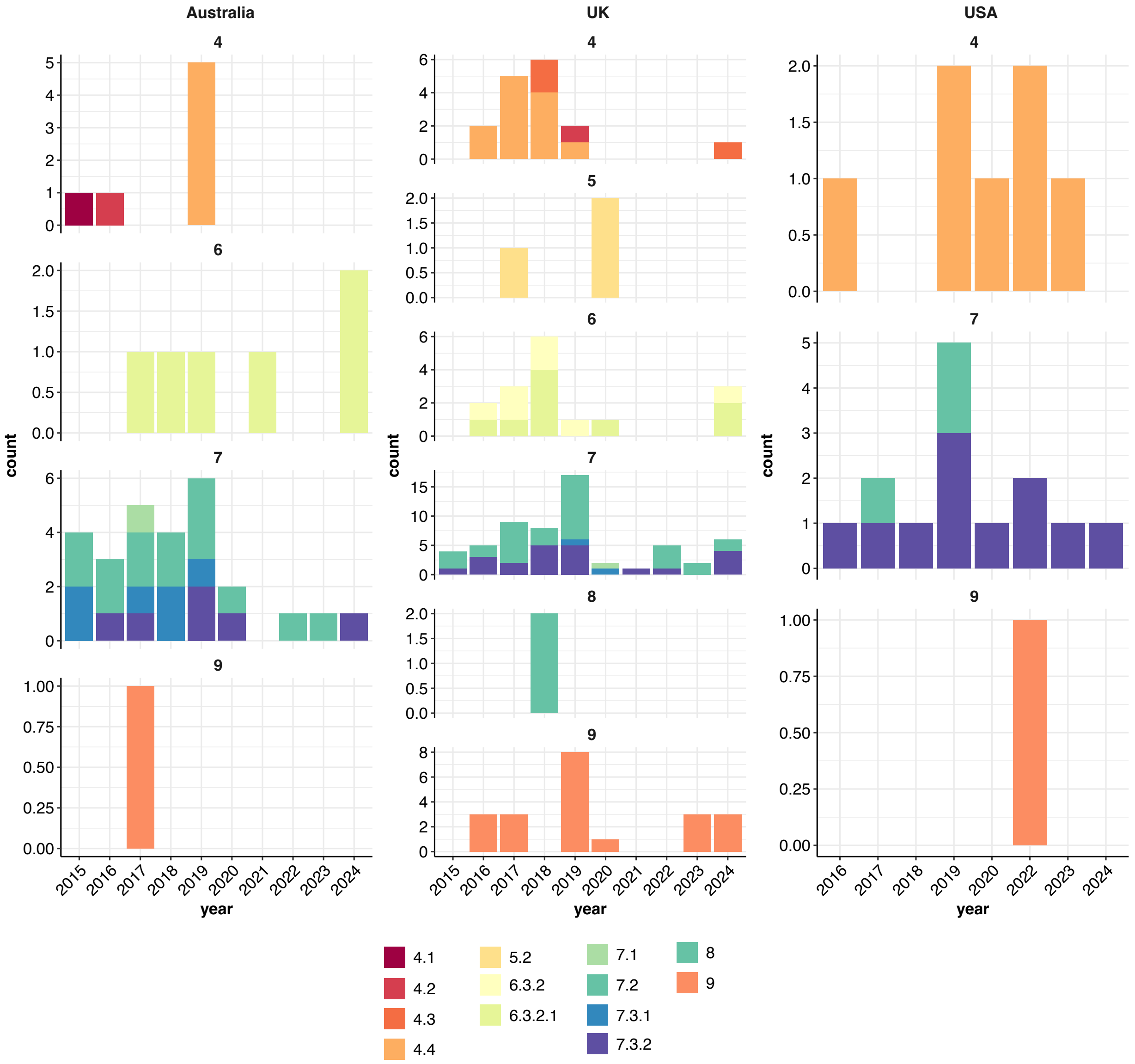


**Fig S4: Barplots of genotypes across three HICs for PGs 4-9**

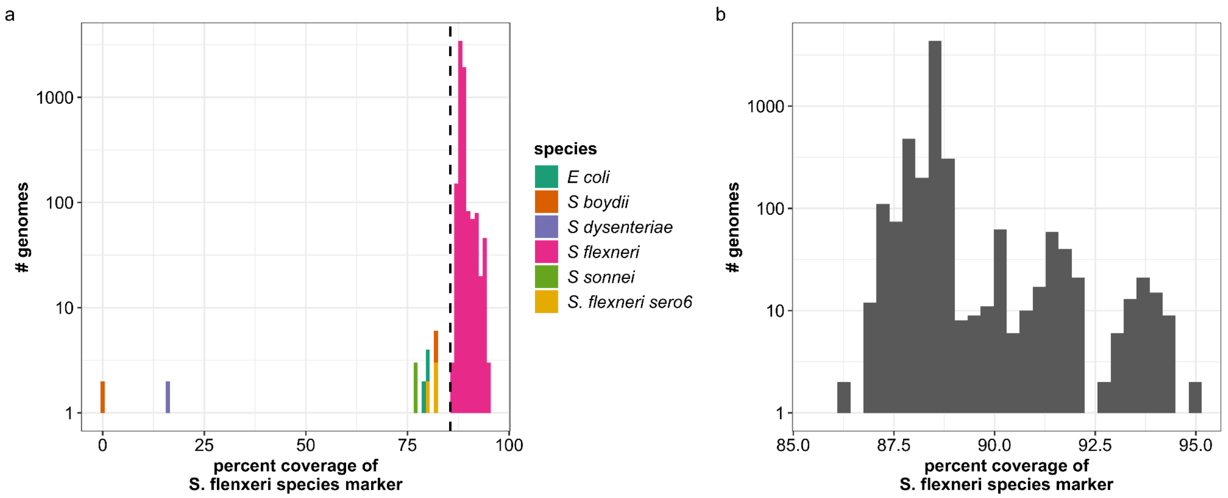


**Fig S5 Species thresholds applied to *S. flexneri,* other *Shigella* species and *E. coli***
A, Distribution of percent coverage to the *S. flexneri* species marker, with bars coloured by actual species. Dashed black line indicates 85.5% coverage which is the threshold for calling as *S. flexneri* for typing purposes. B, Distribution of percent coverage to the species marker for *S. flexneri* genomes only (zoomed in version of pink distribution in panel A).

### List of Supplementary Tables

**Supplementary Table 1:** Details of genomes included in the retrospective dataset
**Supplementary Table 2:** Details of markers in the genotyping scheme
**Supplementary Table 3:** Details of genotype calls from the retrospective dataset that were not a match for the phylogeny
**Supplementary Table 4** Comparison of calls for matched Illumina and ONT data
**Supplementary Table 5:** Details of non *Shigella flexneri* calls using Flex-It
**Supplementary Table 6:** Details of the application of Flex-It to genomes routinely generated in HICs public health laboratories
**Supplementary Table 7:** Coordinates of repeat and phage regions used in masking core SNP alignment

### Additional supplementary files available in FigShare (https://doi.org/10.26180/31697419)

**Supplementary File 1:** Alignment of 5,820 *S. flexneri* isolates (including reference) (.aln)
**Supplementary File 2:** The full discovery and validation tree (.tree)
**Supplementary File 3:** Discovery tree of 2000 isolates (including reference) with nodes labelled (nexus tree)
**Supplementary File 4:** The mutation events in the discovery tree (tsv)
**Supplementary File 5:** The homoplasmic events in the discovery tree (tsv)
